# Molecular mechanisms of LC3-associated phagocytosis in the macrophage response to *Paracoccidioides* spp

**DOI:** 10.1101/2020.10.30.362681

**Authors:** Getúlio Pereira de Oliveira, Herdson Renney de Sousa, Kaio César de Melo Gorgonha, Tatiana Karla dos Santos Borges, Kellyanne Teixeira Rangel, Scott Fabricant, Fernanda Cristina Koser Gustavo, Lucas Fraga Friaça, Angelo Rossi Neto, Fabián Andrés Hurtado, Hugo Costa Paes, Arturo Casadevall, Ildinete Silva-Pereira, Patrícia Albuquerque, Maria Sueli Soares Felipe, André Moraes Nicola

**Author notes:** These authors have contributed equally and share first authorship (Getúlio Pereira de Oliveira, Herdson Renney de Sousa).

## Abstract

Paracoccidiomycosis is a systemic fungal infection that is endemic in Latin America. The etiologic agents are thermodimorphic fungi from the *Paracoccidiodes* genus, which are facultative intracellular parasites of macrophages. LC3-associated phagocytosis (LAP), a noncanonical form of autophagy, is important in the immune response to similar pathogens, so we sought to determine the role LAP plays in the macrophage response to *Paracoccidioides* spp. By immunofluorescence, we found that LC3 was recruited to phagosomes containing *Paracoccidioides* spp. in both RAW264.7 and J774.16 cell lines and in bone marrow-derived macrophages. Interference with autophagy using RNAi against *ATG5* reduced the antifungal activity of J774.16 cells, showing that LC3 recrutiment is important for proper control of the fungus by macrophages. Finally, we used pharmacological Syk kinase and NAPH oxidase inhibitors, which inhibit signalling pathways necessary for macrophage LAP against *Aspergillus fumigatus* and *Candida albicans*, to dissect part of the signaling pathways that trigger LAP agains *Paracoccidioides* spp. Interestingly, these inhibitors did not decrease LAP against *P. brasiliensis*, possibly due to differences in the fungal cell surface compositions. These observations suggest a potential role for autophagy as target for host-directed paracoccidioidomycosis therapies.

## 1. Introduction

The genus *Paracoccidioides* includes several species of thermodimorphic fungi. In nature, it has been isolated from the soil[1] and is commonly found in armadillo burrows[2]. Upon inhalation by humans, *Paracoccidioides* spp. can cause one of the most prevalent systemic mycoses in Latin America – Paracoccidioidomycosis (PCM)[3]. Chronic PCM is characterized by an infection of the respiratory tract, leading to significant morbidity due to lesions in the lung parenchima. Subsequently, the disease can disseminate to other organs and tissues, forming secondary lesions in the mucous membranes, skin, lymph nodes and adrenal glands. Acute PCM, on the other hand, is characterized by disseminated proliferation of fungi in the reticulo-endothelial system, with a high fungal burden in lymph nodes and the spleen[4]. In a hyperendemic area, such as Rondônia state, Brazil, the mean annual incidence was 9.4/100,000 people. The mean annual incidence in a Brazilian state with high incidence was 9.4/100,000 people during 1997-2012, with a case-fatality rate of 10.2% and 9.9% during 2001, and 2002, respectively[5].

The clinical course of PCM is associated with a deficient immune cell response modulated by the balance of cytokines released by inflammatory cells in the microenvironment[6]. The presence of *P. brasiliensis* stimulates monocytes to release pro- and anti-inflammatory cytokines, which trigger an inflammatory granulomatous response characterized by the accumulation of macrophages and effector cells[7]. Macrophages can act in different ways, depending on the activation state of the macrophage. Activated M1 macrophages are pro-inflammatory cells, responsible for phagocytosis and microbe killing. M1 macrophages, activated by IFN-gamma, produce high levels of nitric oxide (NO) and secrete large amounts of IL-12. In contrast, M2 macrophages reside in the tissue and produce high levels of arginase and anti-inflammatory cytokines[8].

PCM treatment usually lasts for six months to two years and many drugs are available. The most commonly used drugs are sulfonamides, amphotericin B and azole derivatives[9]. Treatment is time-consuming and often associated with complications and relapse. Drugs may have undesirable side effects, and some of them are expensive, such as liposomal amphotericin B. Occasional resistant strains have been reported and the search for more selective and efficient antifungals to treat this and other mycoses continues[10]. Thus, the search for molecular targets involved in host protection against fungal invasion may be a viable alternative to the use of antifungal drugs.

Autophagy is an essential process, conserved in all eukaryotes, characterized by the lysosomal degradation of cytoplasmic organelles or cytosolic components[11]. It is activated by nutrient deprivation, hypoxia, oxidative stress, DNA damage, accumulation of protein aggregates or damaged organelles[12]. In addition to endogenous substrates, autophagy is activated in response to intracellular pathogens and can degrade infectious particles and microbes, having a crucial role in resistance to bacterial, viral and protozoan infection in metazoan organisms[13]. These immune functions of autophagy are important against human pathogenic fungi, such as *Aspergillus fumigatus[14], Cryptococcus neoformans* and *Candida albicans[15]*, and in the suppression of immune responses to protect hosts from possible collateral damage caused by overly active immunity[16].

Autophagy requires autophagy-related genes (ATG) for all steps of the process, from phagophore initiation to fusion of the autophagosome with the lysosome. Among autophagy proteins, the Atg8 homolog microtubule-associated protein 1 light chain 3 (LC3) is considered a marker of autophagosome vesicles and participates in a non-canonical autophagy process called LC3 associated phagocytosis (LAP[17]). More recent evidence shows that in the case of fungi, it appears to be LAP, rather than canonical authophagy, that is triggered in response to these agents[18].

Given that the host response to *Paracoccidioides* spp. infection could be targeted for disease prevention or therapy, it is important to understand how mammalian hosts respond to *Paracoccidoides* spp. As LAP has been previously shown to be important in response to other fungi, our aim in this work was to determine the molecular mechanisms involved in LC3 associated phagocytosis in the immune response to *P. brasiliensis*.

## 2. Materials and Methods

### 2.1. Cell lines, fungal strains, and growth conditions

The Pb18 and Pb01 isolates of *P. brasiliensis* and *P. lutzii*, respectively, were maintained in Fava-Netto’s medium (1% w/v peptone, 0.5% w/v yeast extract 0.3% w/v proteose peptone, 0.5% w/v beef extract, 0.5% w/v NaCl, 4% w/v glucose, and 1.4% w/v agar, pH 7.2), at 37 °C. Cultures no older than five days from the last passage were used for experiments. For fungicidal activity experiments, fungal CFUs were counted by plating in brain-heart infusion (BHI) agar supplemented with horse serum and *P. brasiliensis* conditioned medium. All these culture conditions are conducive to the yeast phenotype, which we used for all experiments.

### 2.2. Cell lines

The mouse macrophage cell lines RAW 264.7, and J774.16 were used for the detection of LC3-associated phagocytosis (LAP) *in vitro*. HEK 293T, and J774.16 were used for transfection and transduction assays, respectively. Cells were kept in 100-mm Petri dishes with Dulbecco’s Modified Eagle’s Medium (DMEM), supplemented with non-essential amino-acid solution and 10% of fetal bovine serum (FBS; Thermo Fisher), and incubated at 37 °C and 5% CO_2_.

### 2.3. Animals and primary cells

C57BL/6 mice mice were bred at the Animal Center of the the University of Brasília Institute of Biological Sciences with food and water *ad libitum*. Bone marrow cells from mice six to twelve weeks old were collected. All procedures were performed in accordance with national and institutional guidelines for animal care and were approved by the university’s Institutional Animal Care Use Committee (Proc. UnB Doc 52657/2011). Bone marrow-derived macrophages (BMMs) were generated from bone marrow cells as previously described[19]. Briefly, 2 x 10^6^ bone marrow cells were plated on non-treated 100-mm Petri dishes with RPMI 1640 supplemented with 10% heat-inactivated fetal bovine serum (FBS; Thermo Fisher), 50 μg/mL gentamicin, 50 μM 2-mercaptoethanol (Sigma-Aldrich) and 20 ng/mL recombinant GM-CSF (Peprotech). The cultures were incubated for 8 days at 37 °C in a humidified 5% CO_2_ atmosphere. On day 3, 10 mL of fresh complete medium was added to the culture. Half of the medium was removed at day 6 and new complete medium was added. Attached BMMs were collected on day 8 with TrypLE™ Express (Thermo Fisher).

### 2.4. Production of ATG5 shRNA lentiviral vectors

For the transfection assay we used TAT, REV, GAG-POL and VSV-G vectors engineered from human immunodeficiency virus 1 (HIV-1) and vesicular stomatitis virus (VSV). Plasmid pLK0.1 was used to clone shRNAs for RNA interference with murine *ATG5* as target. Plasmid expansion was performed in thermo-competent *Escherichia coli* (Omnimax T1 cells; Thermo Fisher) in lysogeny broth (LB) with 100 μg/mL ampicillin. Plasmids were purified with the GenElute Plasmid DNA Miniprep Kit (Sigma-Aldrich), quantified using the Qubit fluorometer (Thermo Fisher) and stored at −20 °C.

HEK 293T cells were trypsinized and harvested at 90-95% confluence. Cells were reseeded at a concentration of 3.75 x 10^5^ mL^-1^ onto a six-well plate containing 2 mL of DMEM +10% FBS per well. A mix containing OptiMEM media, Lipofectamine 2000 (Invitrogen), and the assembly media containing the packing plasmids plus the pLKO.1 vectors encoding each shRNA were added to each well. After six hours of incubation, 2 mL of DMEM + 10% FBS were added to the cells, and the supernatant was collected after 12 and 24 h. Supernatants were centrifuged at 200 x g for five minutes to remove dead cells and debris, and the resulting supernatant was centrifuged again at 20.000 x g for 90 min at 4 °C. Pellets containing lentivirus were ressuspended and stored at −80 °C.

### 2.5. J774.16 cell transduction with ATG5 shRNA lentivirus

After J774.16 cells reached 95% confluence, they were harvested by trypsinization and counted. They were reseeded onto a 96-well plate containing DMEM + 10% FBS, at 10^4^ cells/well. In the following day, the cell culture medium was exchanged for DMEM + 10% FBS medium supplemented with 8 μl/mL hexadimethrine bromide (Polybrene), and 5, 50 or 100 μL of lentivirus harbouring each shRNA were added to the cells. In the next day, the medium was exchanged for DMEM + 10% FBS, and after one more day, transduced cells were selected with puromycin at 0.5 μg/mL (Thermo Fisher). After 48 h of selection, untransduced dead cells were removed from the supernatant. For the second round of selection, transduced J774.16 cells were harvested and seeded onto a six-well plate with medium supplemented with puromycin at 5 μg/mL. After reaching confluence, cells were harvested, and total RNA was extracted using Trizol^®^ (Thermo Fisher). RNA was analyzed by electrophoresis with 1% agarose, and gene expression evaluated by qPCR.

### 2.6. Fungal killing assay

For CFU experiments, stably transduced J774.16 cells were seeded onto a 96-well plate at 2 x 10^4^ cells per well in DMEM supplemented with 10% FBS and activated with murine IFN-γ at 200 U/mL plus LPS at 1 μg/mL. After 24 h of activation and adhesion, the J774.16 cells were co-incubated with *P. brasiliensis* suspensions. To prepare these fungal suspensions, *P. brasiliensis* yeast cells were scraped from solid media and suspended in PBS. After vortexing with 2 – 4 mm glass beads for 30 s, large clumps were removed by decanting and the suspension strained through a 40 μm cell strainer. The viable cell density on the resulting suspension was counted in a hemocytometer using the vital dye Phloxine B. J774.16 cells were co-incubated with *P. brasiliensis* for 24 h, with a multiplicity of infection (MOI) of one. After this period, CFUs were counted by plating the same dilution for each well onto BHI agar plates and incubating at 37 °C until colonies appeared (5 to 7 days). Controls included untransduced J774.16 cells and wells with no macrophages. The experiment was repeated independently three times in different days, each with four or five wells per condition.

### 2.7. Co-incubation of macrophages and *Paracoccidioides* spp. for LC3 immunofluorescence

In different experiments, macrophages were either plated onto glass-bottom dishes (Mattek^^®^^) or on 24-well plates with sterile circular coverslips for 24 h. *P. brasiliensis* yeast cells were harvested from five day old culture plates by scraping the surface of the fungal mat, vortexing the cell chunks in PBS, passing the suspension through a 40-μm cell strainer and measuring cell density in a hemocytometer. Fungal cells were inoculated onto the plated macrophages at a MOI of one. The dishes were incubated for 12 to 24 h at 37 °C in the presence of 5% CO_2_ to allow infection. Afterwards, the plates were processed for immunofluorescence as described below. In some experiments, we added the Syk-selective tyrosine kinase inhibitor piceatannol (at 10 or 30 μM) (Invivogen, San Diego, CA, **c**atalog # tlrl-pct) or the NADPH oxidase inhibitor diphenyleneiodonium chloride (DPI; at 10 or 20 μM) (Sigma-Aldrich, Sant Louis, Missouri, catalog #D2926) to the dishes. Inhibitors were added 10 min before stimulation and remained in culture for the duration of the experiments.

### 2.8. LC3 immunolocalization

After 12 or 24 hours of infection, the cells were fixed with ice-cold methanol for ten minutes and washed with PBS. After that, they were incubated with a 1% BSA solution in 1X PBS containing primary antibody (rabbit polyclonal IgG against human LC3, 1:1000 dilution, Santa Cruz Biotechnology) for one hour at 37 °C. They were then washed three times with PBS and incubated with the secondary antibody (goat IgG against rabbit IgG conjugated with AlexaFluor^^®^^ 488, Thermo Fisher Scientific) diluted 1:2000 in the same conditions as the primary one. The cells were afterwards washed three times with PBS and the glass-bottom dishes (or coverslips) were mounted with ProLong Gold Antifade Mountant (Thermo Fisher Scientific). Samples were documented in a Zeiss Axio Observer Z1 epifluorescence microscope equipped with a 63x NA 1.4 oil immersion objective and a cooled CCD camera. Image stacks were deconvolved with a constrained iterative algorithm on Zeiss ZEN and then processed on ImageJ and Adobe Photoshop. No non-linear modifications were made.

### 2.9. Statistical analysis

For *ATG5* knockdown, ANOVA and Dunnett’s multiple comparison pos-hoc tests were performed on Graphpad Prism. For LAP quantitative analysis, a Fisher’s exact test was performed to compare proportions of fungi on LC3-positive vacuoles on Graphpad Prism. For CFU analysis, a mixed-analysis ANOVA was used, with shRNA as a fixed effect and replicate as a random factor. Pairwise comparisons were made using Tukey’s HSD to correct for multiple comparisons. Analysis was performed in R using the MultComp package[20].

## 3. Results

### 3.1. LC3 is recruited to phagosomes containing *Paracoccidioides spp*. in murine macrophages

To test whether macrophages use LAP against *Paracoccidioides* spp., we performed LC3 immunofluorescence experiments with different types of macrophages that had been infected with the fungus. Controls for autofluorescence and non-specific secondary antibody binding were negative (**Figure S1**). LC3 was detected in vacuoles containing *Paracoccidioides* spp. cells in all tested macrophages (RAW264.7, J774.16, and BMM) that had been incubated with *P. brasiliensis* or *P. lutzii* for 12 or 24 hours **(Figure 1A-C)**. These experiments showed that LC3 recruitment does not occur in all vacuoles containing *Paracoccidioides spp*, suggesting that this process might take more time for completion or that macrophages do not use LAP against all internalized fungi. Furthermore, we frequently observed LC3 recruitment around daughter but not mother cells **(Figure 1B)**. Interestingly, we also detected LC3 around apparently extracellular *Paracoccidioides* spp. cells.

**Figure 1 –.**
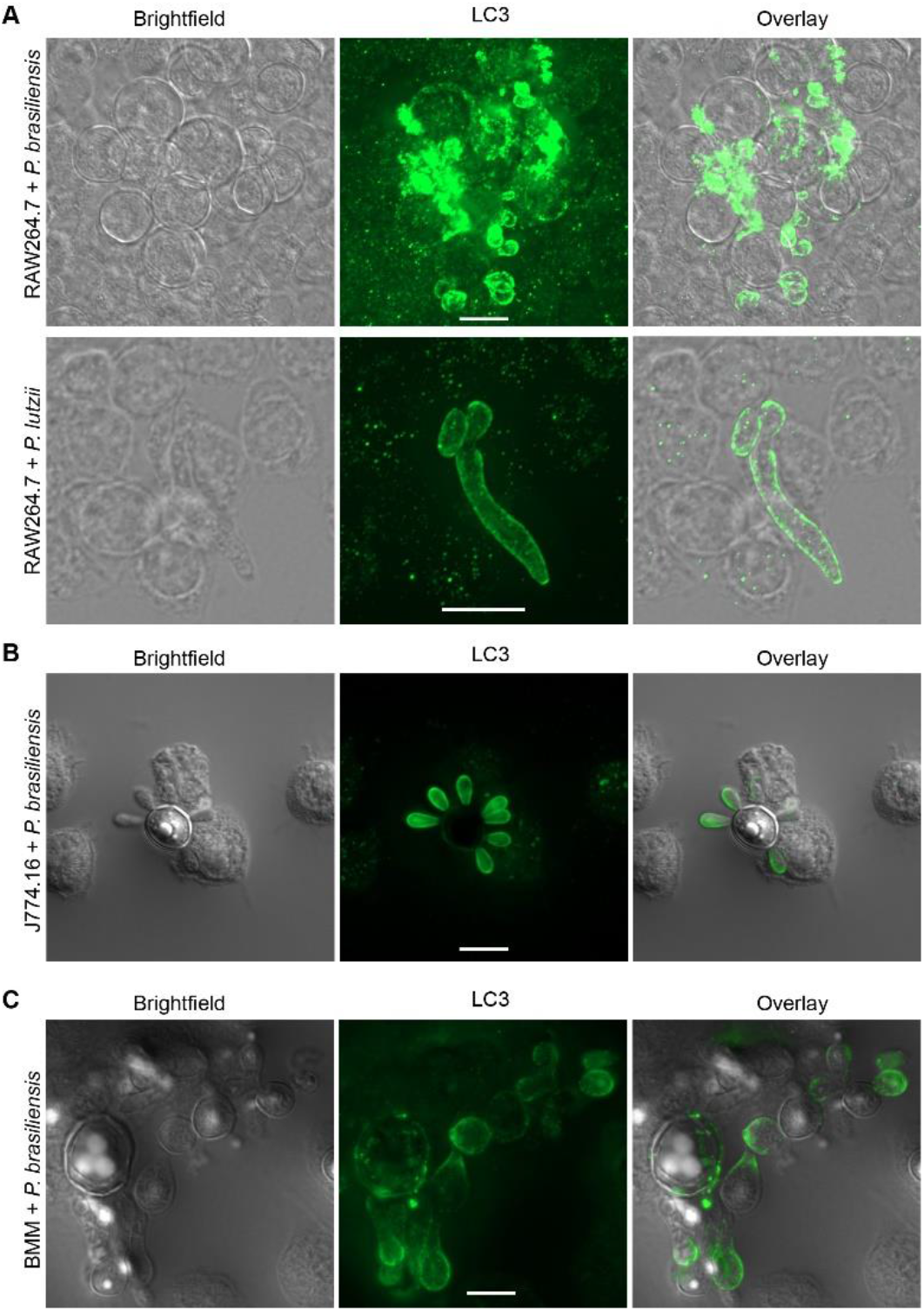
LC3 associated phagocytosis (LAP) is activated in murine macrophages against *Paracoccidioides* spp. **(A)** LAP was detected after 24 h in RAW264.7 incubated with *P. brasiliensis* and *P. lutzii*. **(B)** The same phenomenon was confirmed after 12 h of the *P. brasiliensis* interaction with J774.16 and **(C)** bone marrow derived macrophages (BMM), scale bars: 10 μm. Experiments were repeated at least twice in different days and had similar results.

### 3.2. LAP is important in the murine macrophage response to *P. brasiliensis*

After confirming the occurrence of LAP in murine macrophages incubated with *Paracoccidioides* spp., we performed a loss-of-function experiment by knocking down the *ATG5* gene in J774.16 macrophages. For that purpose, we produced five different lentiviral vectors containing different shRNAs against *ATG5*. J774.16 cells were transduced with lentiviral particles and knockdown levels of *ATG5* were determined by quantitative PCR. The *ATG5* knockdown efficacy varied greatly among the vectors. Two (A, and B) were the most efficient in knocking the gene down, especially in the lowest amount used (5 μL). Vectors A and B reduced *ATG5* gene expression by approximately 97% relative to the negative control EGFP shRNA **(Figure 2A)**. Next, *ATG5*-silenced J774.16 cells were incubated with *P. brasiliensis* and a fungal killing assay (CFU) was performed. When compared to the EGFP control or non-transduced J774.16 cells, the fungal killing assay showed that *ATG5* knockdown significantly increased the survival of *P. brasiliensis* in macrophages **(Figure 2B)**.

**Figure 2 –.**
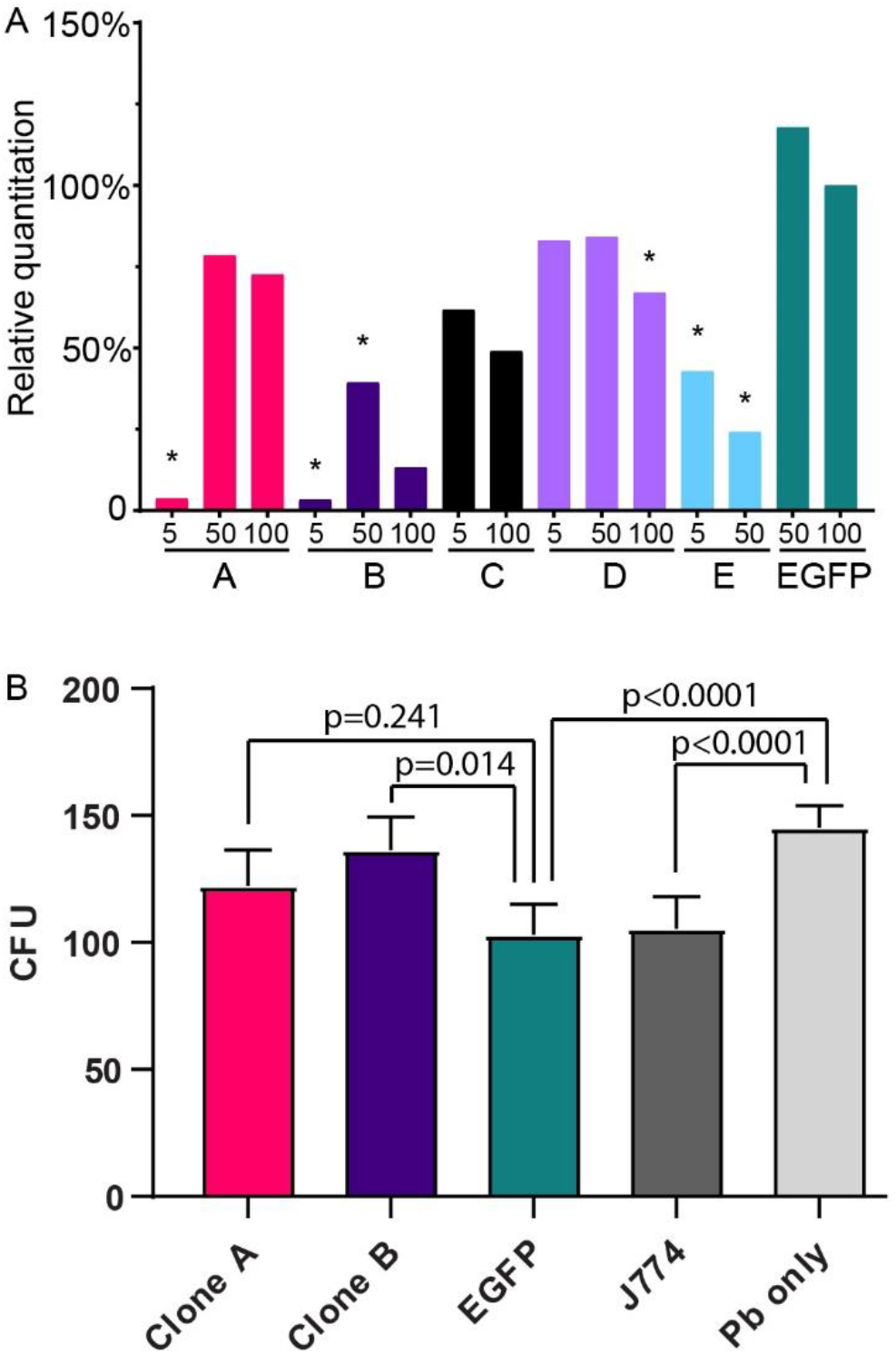
LC3 associated phagocytosis (LAP) is important for the death of *P. brasiliensis* phagocytized by J774.16 murine macrophages. **(A)** Five microlitres of lentiviruses A and B knocked down ATG5 expression by approximately 97% in J774.16 relative to the scrambled EGFP knockdown controls, as measured by real-time RT-PCR. Numbers on the X axis represent the volume of each lentiviral vector used in transduction. * represents p < 0.05 compared to the scramble EGFP (ANOVA and Dunnet multiple comparison pos-hoc test). (B) Fungal killing assay showing that ATG5 knockdown significantly increased the survival of *P. brasiliensis* (Pb) in macrophages, p-value from mixed-analysis ANOVA and Tukey’s HSD multiple comparison pos-hoc test.

### 3.3. Macrophages use a different mechanism to trigger LAP against *Paracoccidioides* spp. in comparison with *C. albicans*

The molecular mechanism of macrophage LAP against *C. albicans* and *Aspergillus fumigatus* has been previously determined[21,22]. Two key steps in this mechanism are dependent on the Syk kinase and the NADPH oxidase complex. To test if the same happened in macrophages infected with *Paracoccidioides* spp., we used pharmacological inhibitors to block these well-described components of autophagy pathways. Surprisingly, the inhibition of Syk by piceatannol led to a dose-dependent increase in LC3-positive phagosomes containing *P. brasiliensis*, whereas inhibition of NADPH oxidase by diphenyleneiodonium chloride (DPI) did not affect LAP (**Figure 3 and Table 1)**. As a control, we used the same inhibitors in macrophages infected with *C. albicans*. As described previously in the literature[21], inhibition of Syk or NADPH oxidase in *C. albicans* leads to a decrease in LAP in BMMs **(Figure S2 and Table 1)**.

**Figure 3 –.**
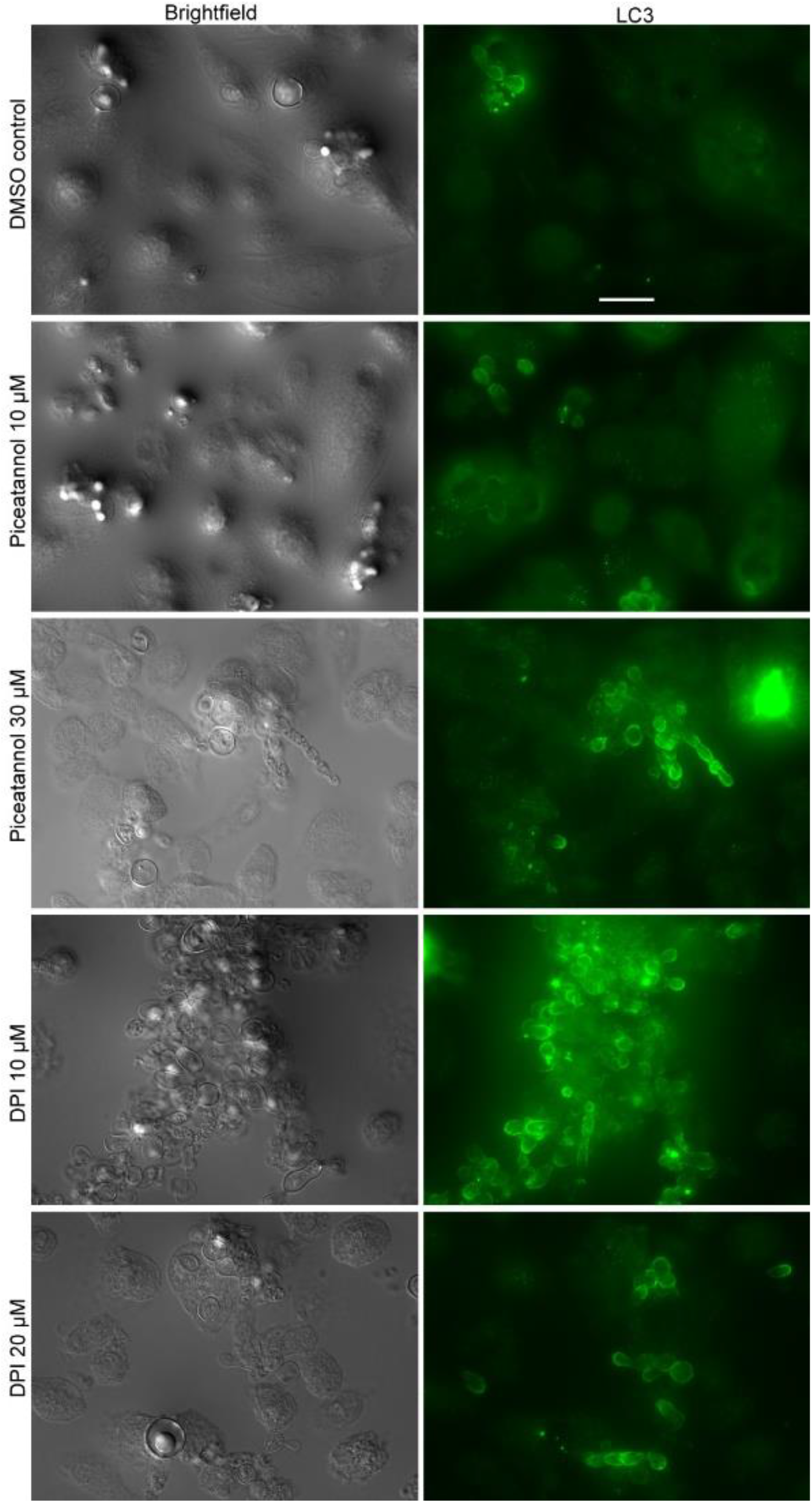
Effect of Syk and NADPH oxidase inhibition on bone marrow derived macrophages (BMM) LAP against *P. brasiliensis*. Piceatannol (10, 30 μM) and diphenyleneiodonium chloride (DPI) (10, 20 μM) were used to inhibit Syk and NADPH oxidase, respectively. Syk inhibition led to a dose-dependent increase in LAP, whereas NADPH oxidase inhibition did not seem to affect LAP. Scale bar: 20 μm.

**Table 1.**
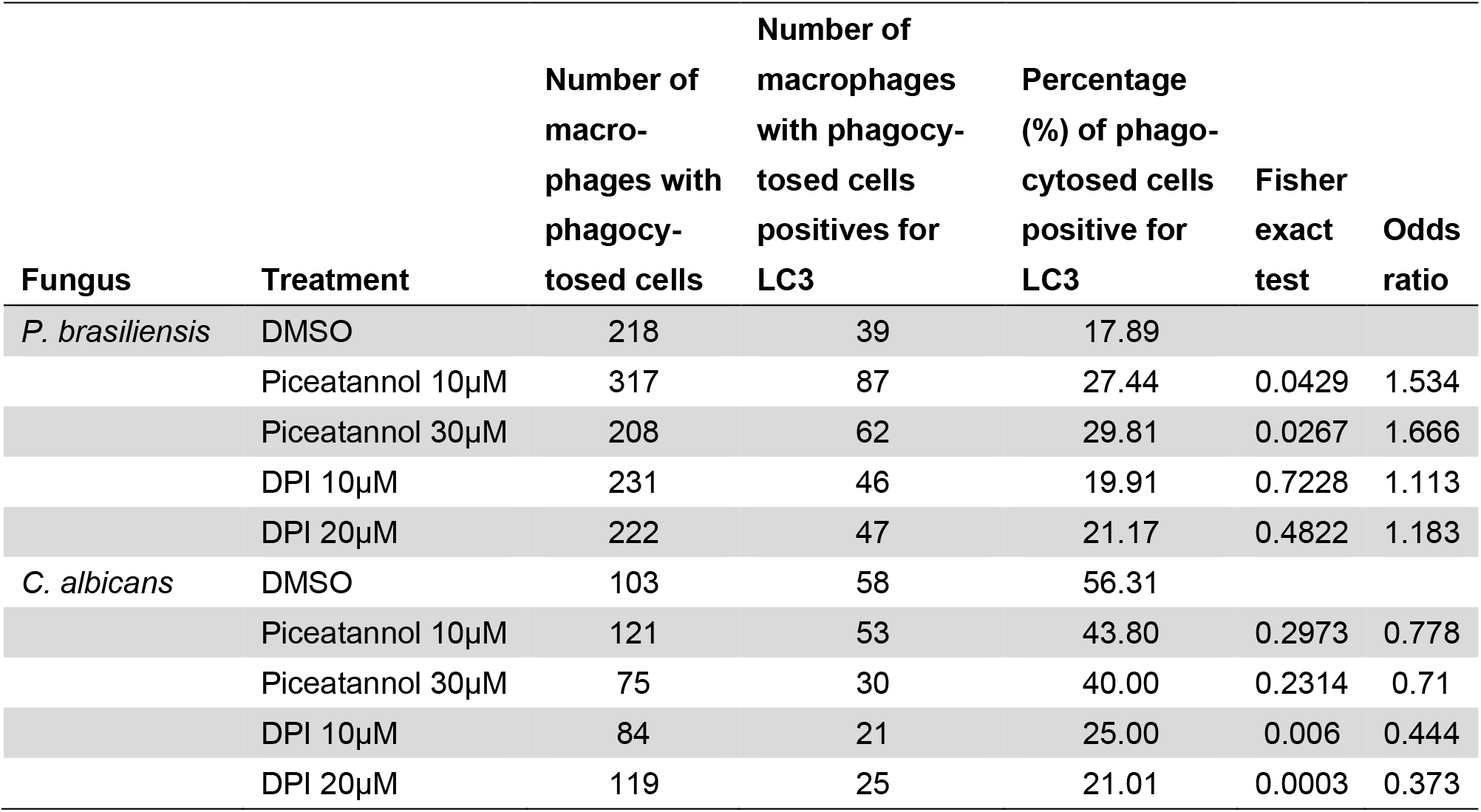
Recruitment of LC3 to vacuoles containing *P. brasiliensis* and *C. albicans* in primary macrophages. BMMs were infected with either fungi in the presence or absence of Syk and NADPH oxidase inhibitors and processed for immunofluorescence microscopy as shown in Figure 3. The total number of ingested fungi and the number of fungi on LC3-positive vacuoles was then counted.

## 4. Discussion

About 1.5 million people die every year from systemic fungal infections[23]. Most of these diseases can only be treated with a small number of drugs, which are often toxic, expensive or take a long time to be effective. This is especially true for paracoccidioidomycosis, which in less severe cases are usually treated with sulfonamides for 12 – 24 months and in more severe cases is treated with the nephrotoxic amphotericin B or its expensive lipid formulations[24]. Even when therapy successfully clears the fungal infection, as many as 40% of the patients have fibrotic sequellae as a result of pulmonary inflammation[25,26]. Thus, host-targeted therapies that modulate the antifungal immune response have a great potential in the therapy of PCM and other systemic mycoses. Our results suggest that LAP might be one such possible target for host-directed therapies.

LAP is a form of non-canonical autophagy and plays important roles in the macrophage immune response against microbes[27], including fungi[28]. It has been studied in the macrophage response to *Histoplasma capsulatum*[29,30], *C. albicans[15,31], C. neoformans[15,32], Aspergillus fumigatus*[14,22] and *Saccharomyces cerevisiae*-derived zymosan[33]. Our results show that LAP is also used by different types of immortalized and primary murine macrophages against two species of the genus *Paracoccidioides*. LC3 accumulated differently around buds and mother cells and was only found in part of the phagosomes containing *Paracoccidioides* spp. cells, which could suggest these fungi might evade LAP. Such immune evasion has been observed in macrophages interacting with *A. fumigatus*, which use an outer layer of melanin to shield cell wall PAMPs from recognition by receptors that trigger LAP[14].

This interpretation is also compatible with our findings with Syk kinase and NADPH oxidase inhibitors. In macrophages that have ingested *C. albicans*[31], *A. fumigatus*[22] or *H. capsulatum*[29], LC3 recruitment to phagosomes initiates after Dectin-1 recognition of β-glucans, followed by Syk activation and NADPH oxidase-dependent generation of reactive oxygen species. On the other hand, LAP has been found to be triggered by other pattern recognition receptors such as TLR2 [33] or even phagocytic receptors such as those for Fcγ[34] or complement[35]. These different LAP pathways may explain our observation of the opposing LAP effects of Syk and NADPH inhibitors in macrophages challenged with *P. brasiliensis* or *C. albicans*.

Our fungal killing assays with ATG5 shRNAs suggest the recruitment of LC3 to phagosomes containing *Paracoccidioides* spp. seems to play a role in their proper antifungal activity. The absolute differences in CFUs between control and ATG5-knockdown cells were not very large (16.8% for clone A and 29.7% for clone B, in comparison with the EGFP control). However, these small differences might be due to the overall limited antifungal effect of J774.16 cells, as indicated by the fact that the EGFP control cells themselves only reduced the CFU counts by 31.3% in comparison with the wells that contained fungi without macrophages. This host-protective role of autophagy *in vitro* is consistent with results our group has obtained in an unrelated project (manuscript submitted). In that work, we tested if there were differences in LAP between dendritic cells obtained from two mouse strains, one of which resistant and the other susceptible to *P. brasiliensis* infection. The percentage of LC3-positive phagosomes was higher in the resistant strain, which constitutes indirect evidence that LAP might play a role in murine infections.

Despite the congruent evidence from our experiments *in vitro* and those with macrophages and dendritic cells from susceptible and resistant mouse strains our group has done (manuscript submitted), care is warranted in reaching strong conclusions regarding macrophage LAP in immunity to *Paracoccidioides* spp. In macrophages infected with the closely related *H. capsulatum*, LAP is actually detrimental to the host and exploited by the fungus to survive[30]. Moreover, the literature on antifungal LAP in macrophages is ripe with apparent contradictions that highlight how complex this mechanism is. In macrophages infected with *C. neoformans* in vitro, for instance, we found that LAP was host-protective[15] but another group found it benefitted the pathogen[32,36]. In invasive candidiasis models, we[15] and others[16,21,37] found that autophagy was host-protective, whereas other experiments showed it was not necessary for proper responses to *C. albicans*[38]. As further experiments with *Paracoccidioides* spp. and other fungi uncover more details on the mechanism that triggers LAP and its role on immune effector functions, we might be able to rely on a new generation of specific autophagy modulating compounds[39–42] for host-directed therapy in paracoccidioidomycosis.

## Author Contributions

Conceptualization: A.M.N., A.C. and P.A.; writing—original draft preparation: G.P.O.J., H.R.S., K.C.M.G. and A.M.N; supervision: A.M.N., I.S.P. and M.S.S.F.; cell lines, and fungal strains maintenance: H.C.P., I.S.P., P.A., M.S.S.F. and A.M.N.; production of ATG5 shRNA lentiviral vectors, transduction, transfection, and fungal killing assay: K.C.M.G., T.K.S.B., K.T.R., S.F., F.C.K.G., L.F.F., A.R.N. and H.C.P.; BMM production: F.A.H.; co-incubation of macrophages and *Paracoccidioides* spp. for LC3 immunofluorescence: G.P.O.J. and H.R.S.; LC3 immunolocalization: G.P.O.J., H.R.S., K.T.R. and F.C.K.G.; Statistical analysis of the data: G.P.O.J., H.R.S., S.F. and A.M.N.; Review: H.C.P., I.S.P., P.A. and M.S.S.F.; Funding acquisition: A.M.N., I.S.P., A.C. and M.S.S.F. All authors have read and agreed to the published version of the manuscript.

## Funding

A.M.N was funded by FAP-DF awards 0193.001048/2015-0193.001561/2017 and the CNPq grant 437484/2018-1. M.S.S.F was supported by FAP-DF/PRONEX award 193.001.533/2016. G.P.J. was supported by a scholarship from Capes – number 150510/2017-9.

## Acknowledgments

The authors would like to thank John Reidhaar-Olson, from the Albert Einstein College of Medicine shRNA Core Facility, for the assistance with shRNA experiments.

## Conflicts of Interest

The authors declare that they have no conflicts of interest.

## Supplementary Material

**Figure S1 –.**
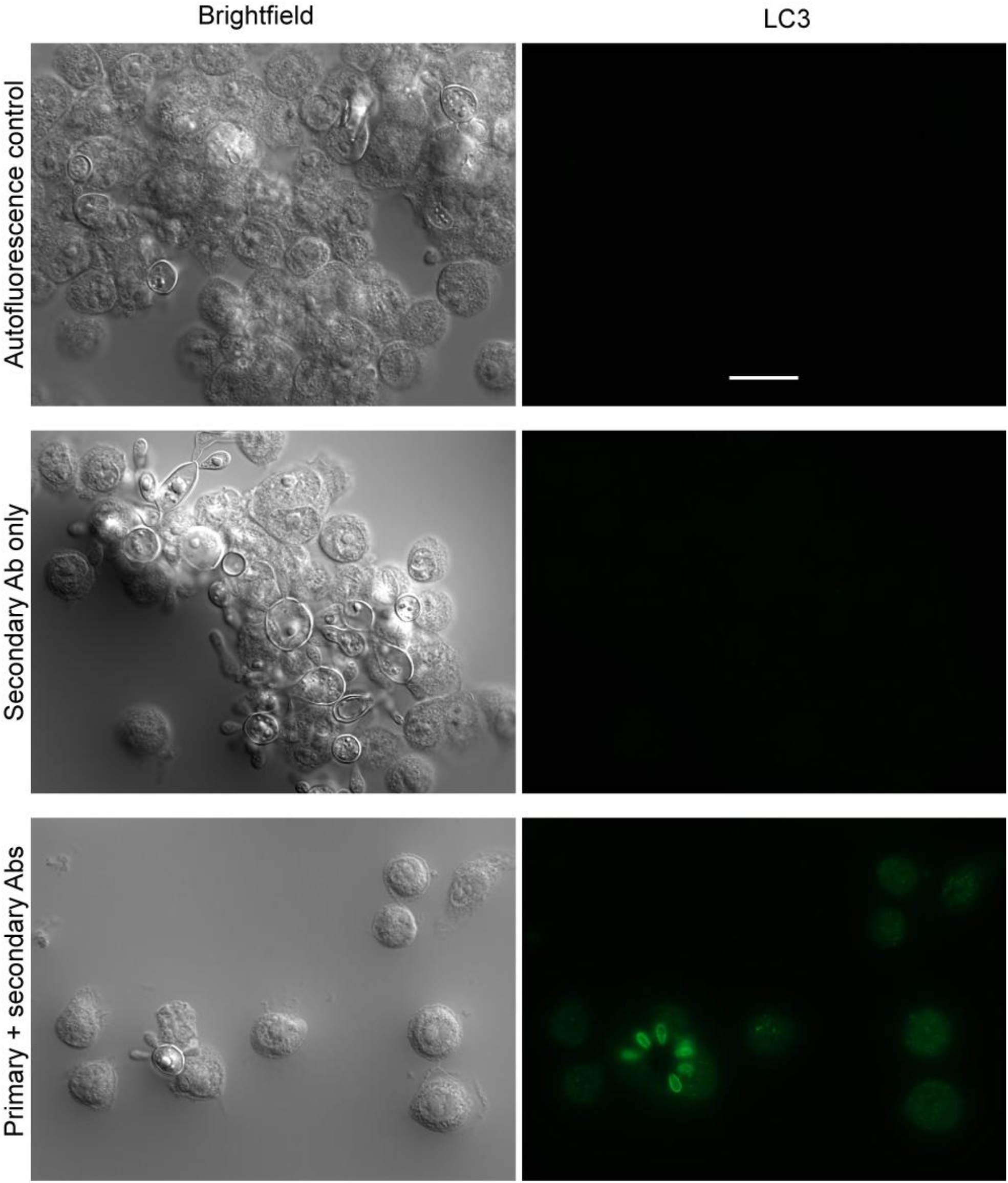
Immunofluorescence microscopy controls. Autofluorescence and non-specific secondary antibody binding controls were used to verify our LAP experiments. No fluorescence was identified in either control experiments. The bottom panel is the whole field of figure 1B. Scale bar: 10 μm.

**Figure S2 –.**
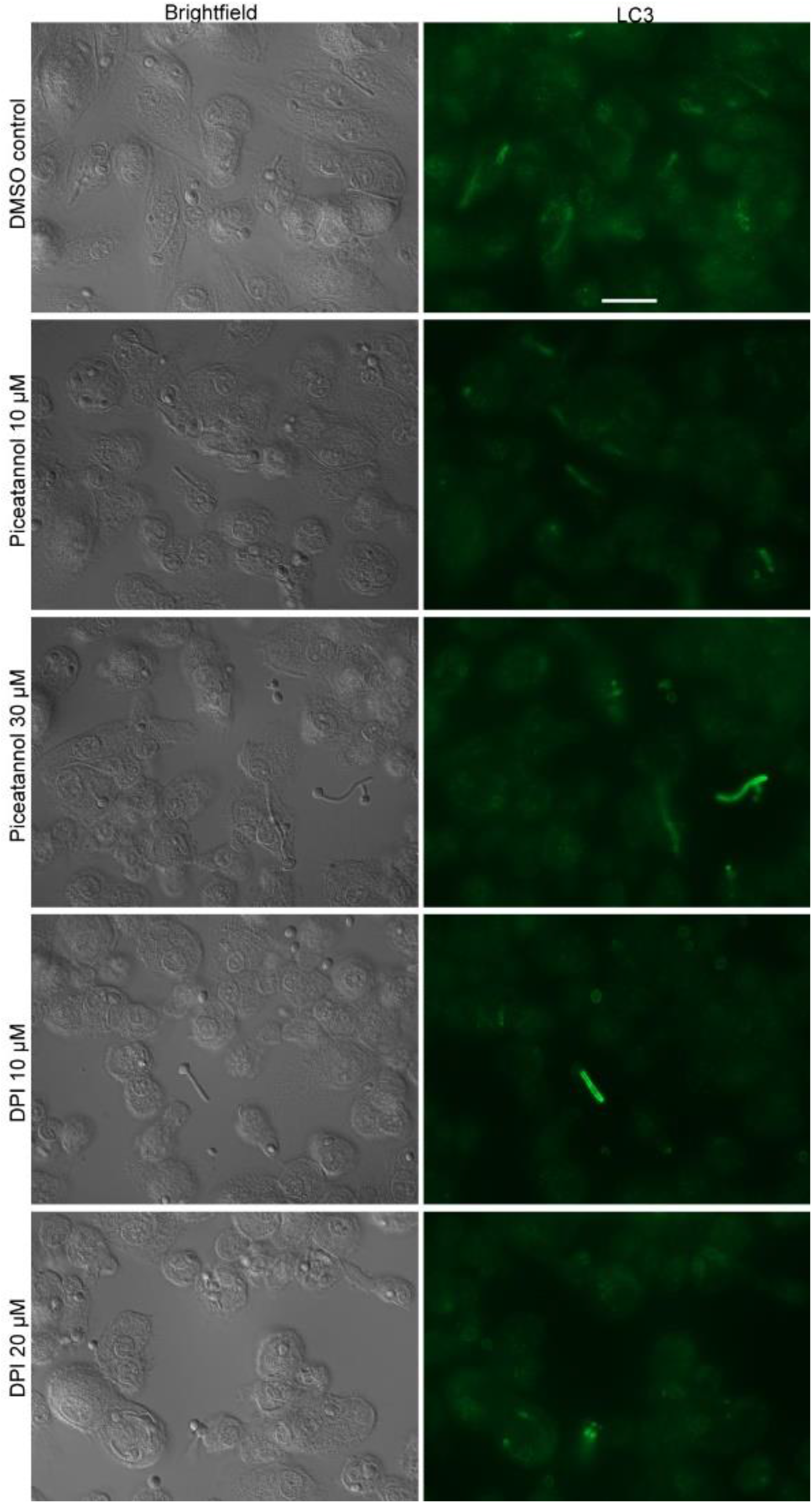
Inhibition of Syk and NADPH in macrophages infected with *C. albicans*. BMMs were infected with *C. albicans* for 12 h in the presence of Syk and NADPH oxidase inhibitors. The cells were then used for LC3 immunofluorescence, showing a decrease in LAP. Scale bar: 20 μm

## Notes

### Competing Interest Statement

The authors have declared no competing interest.

